# Vascular Endothelial Cells Perform Distinct Sensing and Signaling of Laminar and Disturbed Flows across Plasma Membranes and Mitochondria

**DOI:** 10.1101/2023.04.20.537745

**Authors:** Kimiko Yamamoto, Ryohei Maeno, Kenshiroh Kawabe, Yuji Shimogonya, Joji Ando

## Abstract

**BACKGROUND:** Vascular endothelial cells (ECs) experience two different blood flow patterns: laminar and disturbed flows. Their responses to laminar flow contribute to vascular homeostasis, whereas their responses to disturbed flow result in EC dysfunction and vascular diseases. However, it remains unclear how ECs differentially sense laminar and disturbed flows and trigger signalings that elicit different EC responses. We aimed to investigate EC flow-sensing and signaling mechanisms, focusing on the role of the plasma membrane and mitochondria.

**METHODS:** We exposed cultured human aortic ECs to laminar flow and disturbed flow in flow-loading devices and used real-time imaging with optical probes to examine changes in the lipid order of the plasma and mitochondria membranes and the mitochondrial adenosine triphosphate (ATP) production and hydrogen peroxide (H_2_O_2_) release.

**RESULTS:** The lipid order of EC plasma membranes immediately decreased in response to laminar flow, while it increased in response to disturbed flow. Laminar flow also decreased the lipid order of mitochondrial membranes and increased mitochondrial ATP production. In contrast, disturbed flow increased the lipid order of mitochondrial membranes and increased the release of H_2_O_2_ from mitochondria. Addition of cholesterol to the cells increased the lipid order of both membranes and abrogated the laminar flow-induced ATP production, while treatment of the cells with a cholesterol-depleting reagent, methyl-β cyclodextrin, decreased the lipid order of both membranes and abolished the disturbed flow-induced H_2_O_2_ release, indicating that the changes in the membrane lipid order are closely linked to the flow-induced changes in the mitochondrial functions.

**CONCLUSIONS:** ECs differentially sense laminar and disturbed flows by altering the lipid order of their plasma and mitochondrial membranes in opposite directions, which result in distinct changes in the mitochondrial functions, namely, increased ATP production for laminar flow and increased H_2_O_2_ release for disturbed flow, leading to ATP- and H_2_O_2_-mediated signalings, respectively.

## Novelty and Significance

### What is known?

- Vascular endothelial cells (ECs) respond differently to laminar and disturbed flows through undergoing differential alterations of their morphology, functions, and gene expressions.
- The EC plasma membrane acts as a mechanosensor to sense laminar flow through undergoing rapid changes in its physical properties, such as the lipid order and membrane fluidity.
- The EC mitochondria function as signaling organelles that transmit information into the cells when the cells are subjected to changes in their environmental conditions.

### What New Information Does This Article Contribute?

- ECs differentially sense laminar and disturbed flows through undergoing opposite changes in the lipid order of their plasma membrane, triggering flow-specific signaling via mitochondrial responses.
- Upon exposure of ECs to laminar and turbulent flows, the lipid order of not only the plasma membranes, but also of the mitochondrial membranes undergoes changes in opposite directions.
- The EC show intracellular signaling specific for each type of flow through different mitochondrial responses: the EC mitochondria respond to laminar flow by increased ATP production, and to disturbed flow by increased H_2_O_2_ release.

How ECs sense the two blood flow patterns, namely, laminar and disturbed flows, and trigger differential cell signalings still remain unclear. Here, we show that ECs sense laminar and disturbed flows through undergoing differential changes in the lipid order of their plasma membranes, with exposure of the cells to laminar flow decreasing, and exposure to disturbed flow increasing the lipid order. Similar lipid order changes occur almost simultaneously in the mitochondrial membranes, which are then linked to changes in the mitochondrial functions (i.e., increased ATP production for laminar flow and increased H_2_O_2_ release for disturbed flow). Thus, ECs seem to have distinct sensing and signaling mechanisms for these two flow patterns, involving both the plasma membrane and mitochondrial membrane/functions, leading to ATP-mediated purinergic signaling for laminar flow and H_2_O_2_-mediated redox signaling for disturbed flow.

Vascular endothelial cells (ECs) sense shear stress generated by flowing blood and transmit this information into the cell interior, leading to EC responses involving changes in morphology, function, and gene expression.^1^ ECs can experience two different flow patterns depending on their location in the vascular trees. Unidirectional laminar flow occurs in straight conduit arteries, whereas disturbed flow with unsteady direction and velocity occurs in certain areas of arterial bends and branches. Many studies have shown that ECs respond differently to laminar flow and disturbed flow, and that EC responses to laminar flow contribute to the maintenance of vascular homeostasis, while responses to disturbed flow lead to impaired EC function and the development of vascular diseases such as aneurysms and atherosclerosis.^2–4^ However, it remains unclear how ECs separately sense laminar and disturbed flows and transmit this information into the cell interior to elicit different EC responses.

EC sensing and signaling of shear stress has been studied extensively, with a focus on laminar flow. Laminar flow has been shown to activate a wide variety of intracellular signaling pathways via membrane molecules, such as ion channels, receptors, adhesion molecules and proteoglycans, cytoskeletons, and membrane microdomains, including caveolae and primary cilia; these pathways resulted in changes in various EC functions.^5^ Furthermore, the plasma membrane itself has been shown to play an important role in the sensing and signaling of laminar flow.^6^ EC plasma membranes rapidly respond to laminar flow by altering their physical properties, such as their fluidity and viscosity, leading to the activation of G proteins and mitogen-activated protein kinase.^7–10^ Our previous study showed that laminar flow immediately altered the physical properties of the EC plasma membrane by decreasing the lipid order, increasing membrane fluidity, and decreasing the membrane cholesterol content, which triggered ATP release into the extracellular space and purinergic-receptor-mediated Ca^2+^ signaling.^11–13^ This Ca^2+^ signaling plays a crucial role in the control of vascular tone and remodeling through the production of a potent vasodilator, nitric oxide.^14^ Thus, the sensing and signaling of laminar flow has been elucidated to a large extent; that of disturbed flow, however, remains largely unknown.

For some time now, mitochondria have been attracting attention for their function as signaling organelles that transmit information into the cell when the cells are subjected to changes in environmental conditions.^15–17^ Recent data have shown a direct role of mitochondria in the EC mechanotransduction of fluid shear stress.^18, 19^ We previously demonstrated that the application of laminar flow to ECs immediately increased mitochondrial ATP production by enhancing oxidative phosphorylation, leading to the above-mentioned ATP release and purinergic Ca^2+^ signaling.^13, 20^ However, it is not yet known how shear stress acting on the plasma membrane alters intracellular mitochondrial function. Recently, it has become apparent that mitochondrial functions are regulated not only by the concentration of oxygen and ADP in the matrix and mitochondrial membrane potential, but also by the physical properties of the mitochondrial membrane, such as viscosity and lipid order.^21^ For example, oxidative phosphorylation was shown to be enhanced when the viscosity of the mitochondria inner membrane was decreased by increasing the amount of unsaturated lipids. However, whether the flow stimulation of ECs affects the physical properties of mitochondrial membranes has not been investigated to date. The inability to measure the physical properties of mitochondrial membranes has prevented such research, but a new imaging method has recently been developed, making this possible.^22^

The present study aimed to elucidate how ECs perform sensing and signaling for laminar flow and disturbed flow, focusing on the roles of the plasma membrane and mitochondria. To achieve this purpose, we applied laminar flow or disturbed flow to cultured human aortic ECs (HAECs) in flow-loading devices and analyzed the changes in the lipid order of both plasma membranes and mitochondrial membranes and in mitochondrial ATP production and hydrogen peroxide (H_2_O_2_) release using real-time imaging methods with various optical probes.

## Results

### Laminar flow rapidly decreased the lipid order of plasma membranes, whereas disturbed flow increased the lipid order

HAECs were subjected to laminar or disturbed flow in flow-loading devices and the resulting changes in the lipid order of their plasma membranes were examined using real-time Laurdan imaging. To apply disturbed flow to cells, we used a parallel-plate flow chamber in which a step was placed upstream of the flow path; disturbed flow was then generated behind the step (Figure 1E). Normalized transverse wall shear stress (NtransWSS), an indicator of the degree of disturbance in the strength and direction of the shear stress, was calculated using a computer fluid dynamics (CFD) analysis, and cells subjected to a disturbed flow of 0.2 or more NtransWSS (average, 0.4) were analyzed. On the other hand, a parallel-plate flow chamber with no step was used to apply laminar flow with zero NtransWSS to cells.

**Figure 1.**
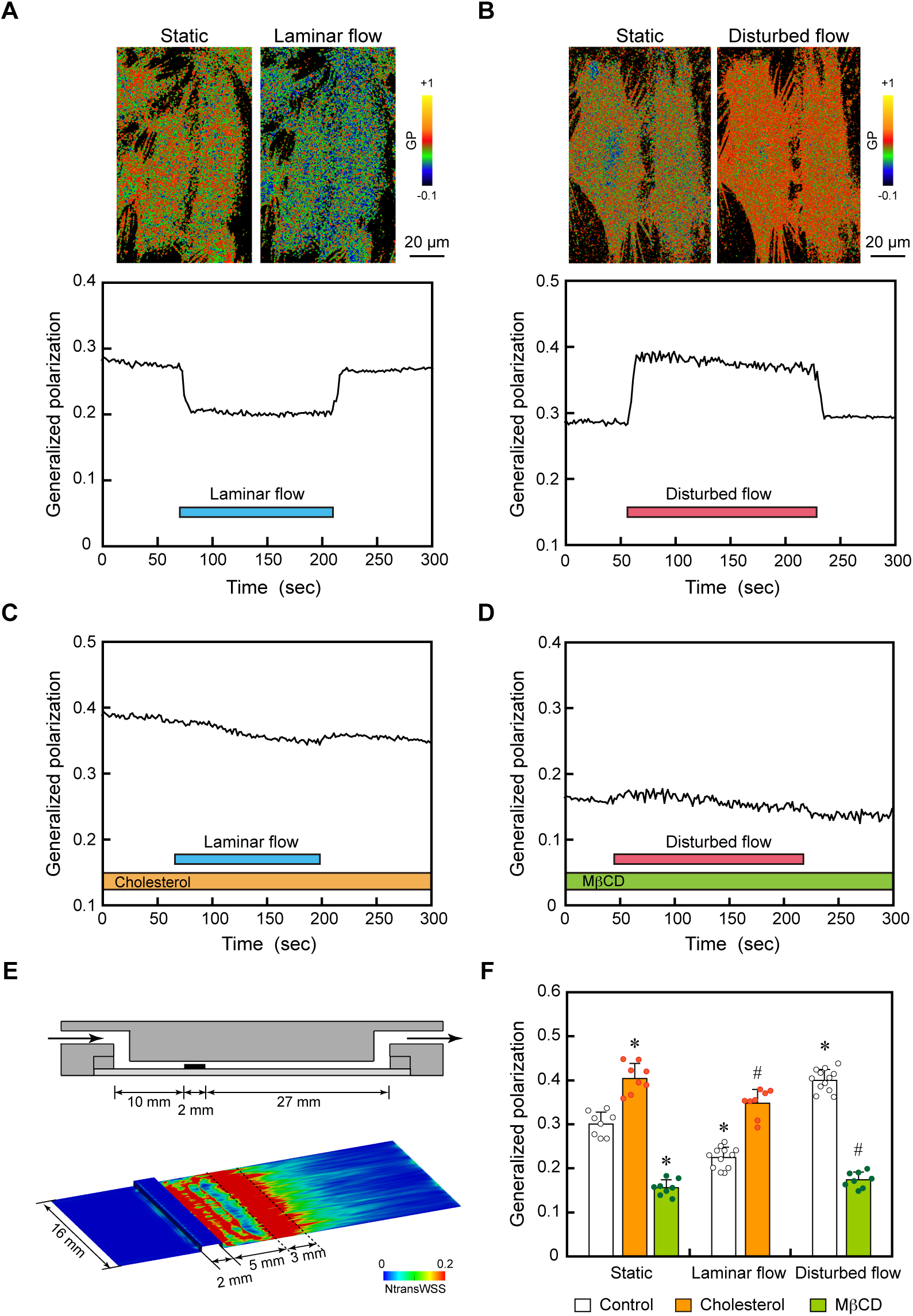
Laminar versus disturbed flow effects on the lipid order of plasma membranes. (**A**, **B**) 3D-reconstructed Laurdan generalized polarization (GP) pseudocolor images of human aortic ECs (HAECs) before and 2 min after laminar flow or disturbed flow. The colors represent different GP values, an indicator of membrane lipid order, according to the scale. The pseudocolor images and temporal changes in GP values showed that the EC membrane lipid order clearly decreased in response to laminar flow, while it clearly increased in response to disturbed flow. (**C**) Addition of cholesterol (100 μM) to ECs prevented the laminar flow-induced decrease in plasma membrane lipid order. (**D**) Treatment of ECs with methyl-β-cyclodextrin (MβCD; 10 mM), a membrane cholesterol depleting agent, abolished the disturbed flow-induced increase in plasma membrane lipid order. (**E**) Flow-loading devices and CFD analysis. In the parallel-plate flow chamber with a step, cells in the range of 5-8 mm downstream of the step (in red) were subjected to disturbed flow with an NtransWSS of greater than 0.2 (0.4 on average) and an average WSS of 20 dyne/cm^2^. On the other hand, in the flow chamber without a step, cells were subjected to laminar flow with an NtransWSS of zero and a WSS of 15 dyne/cm^2^. (**F**) Quantitative analysis of the changes in the GP values induced by laminar flow and disturbed flow, with or without the addition of cholesterol and treatment with MβCD. Values are the means ± SD of data obtained from the samples shown as points. \**P* < 0.01 vs. static control. ^#^*P* < 0.01 vs. each flow control.

Images of Laurdan-labeled HAECs were obtained using a confocal microscope equipped with a two-photon laser, and images of generalized polarization (GP), an indicator of membrane lipid order, were reconstructed as described in the Materials and Methods section. The GP images shown in pseudocolor demonstrated a heterogeneous distribution of GP values across the cell surface, indicating that EC membranes have a nonuniform lipid order in which both liquid-ordered states with high GP values and liquid-disordered states with low GP values coexist (Figure 1A, B).

When laminar flow was applied to cells, the GP images showed a decrease in the lipid order over the entire plasma membrane (Figure 1A). The temporal changes in lipid order were quantified by placing regions of interest (ROIs) throughout the cell. The GP values decreased right after the onset of flow, plateaued thereafter, and returned to the control level after the flow ceased. The laminar-flow-induced decrease in membrane lipid order was similar to what we previously observed in human pulmonary artery ECs.^11^ In contrast, when the cells were subjected to disturbed flow, the GP images showed an increase in the lipid order over the entire plasma membrane (Figure 1B). The GP values began to increase immediately after the application of disturbed flow, remained at an increased level, and then returned to the control level after the flow ceased. The data obtained from many cells confirmed that EC membranes exhibit contrasting responses to laminar flow and disturbed flow by changing their membrane lipid order in opposite directions (Figure 1F). This indicates that EC plasma membranes distinguish between the two different flow patterns through their lipid order responses.

The addition of cholesterol to the cells increased the lipid order of plasma membranes under static conditions and abolished the lipid order lowering effect of laminar flow (Figure1C, F). On the other hand, the treatment of ECs with methyl-β-cyclodextrin (MβCD), which extracts membrane cholesterol, reduced the lipid order of plasma membranes under static conditions and markedly prevented the lipid order-increasing effect of disturbed flow (Figure 1D, F). These findings indicate that adding or removing cholesterol to cells directly alters the lipid order of plasma membranes and can prevent the responses of the membrane lipid order to laminar and disturbed flows.

### Laminar flow rapidly decreased the lipid order of mitochondrial membranes, whereas disturbed flow increased the lipid order

HAECs were exposed to laminar flow or disturbed flow, and changes in the lipid order of the mitochondrial membranes were examined using real-time imaging with the solvatochromic fluorescent probe NRMito, a Nile Red derivative bearing chemical groups that target mitochondria specifically.^22^ The localization of NRMito in HAECs coincided precisely with the mitochondria, as shown by live-cell imaging using the mitochondrion-selective dye MitoTracker (Figure 2A). Ratiometric images of NRMito-labeled HAECs were reconstructed as described in the Materials and Methods section and displayed in pseudocolor.

**Figure 2.**
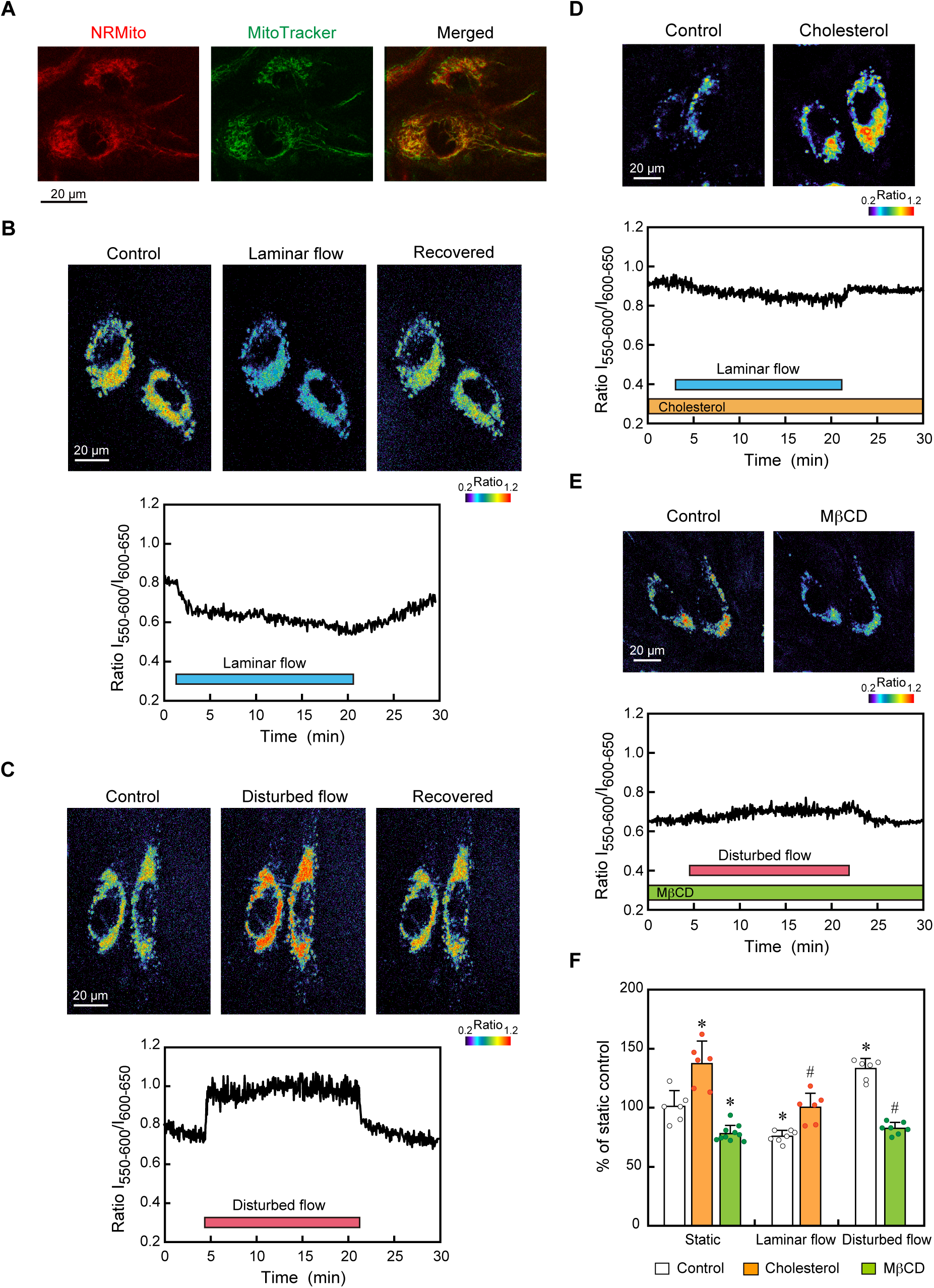
Laminar versus disturbed flow effects on the lipid order of mitochondrial membranes. (**A**) Confocal images of live HAECs expressing the solvatochromic probe NRMito (in red) and labeled with the mitochondrion selective dye MitoTracker (in green). NRMito was correctly located in mitochondria. (**B**) Pseudocolor images of the NRMito fluorescence intensity ratio obtained before, during, and after the application of laminar flow. These images and the temporal change in the NRMito ratio showed that the mitochondrial membrane lipid order rapidly decreased in response to laminar flow. (**C**) The pseudocolor images and the temporal change in the NRMito ratio showed that the lipid order of the mitochondrial membranes rapidly increased in response to disturbed flow. (**D**) Effects of cholesterol addition to ECs on the mitochondrial membrane lipid order and its response to laminar flow. The pseudocolor images obtained 6 h after the addition of cholesterol (100 μM) showed an increase in the lipid order of the mitochondrial membranes. The temporal change in the NRMito ratio showed that cholesterol addition prevented the laminar flow-induced decrease in the lipid order. (**E**) Effects of cholesterol extraction by treating ECs with MβCD (10 mM) on the mitochondrial membrane lipid order and its response to disturbed flow. The pseudocolor images obtained 30 min after MβCD treatment showed a decrease in the lipid order of the mitochondrial membrane. The temporal change in the NRMito ratio showed that cholesterol extraction abolished the disturbed flow-induced increase in the lipid order. (**F**) Quantitative analysis of the changes in the NRMito ratio induced by laminar flow and disturbed flow, with or without the addition of cholesterol and treatment with MβCD. Values are the means ± SD of the data obtained from samples shown as points. \**P* < 0.01 vs. control. ^#^*P* < 0.01 vs. each flow control.

Pseudocolor images showed that laminar flow reversibly decreased the NRMito ratio, i.e., the lipid order of the mitochondrial membrane (Figure 2B). As shown in the temporal change of the NRMito ratio, the lipid order decreased right after the onset of laminar flow and plateaued thereafter; the decreased level returned to the control level after the cessation of flow. In contrast, pseudocolor images demonstrated that disturbed flow reversibly increased the lipid order of the mitochondrial membrane (Figure 2C). The temporal change showed that the lipid order began to increase immediately after the application of disturbed flow, remained at an increased level, and then returned to the control level after the flow ceased. The data obtained from many cells confirmed that mitochondrial membranes exhibit contrasting responses to laminar flow and disturbed flow by changing their membrane lipid order in opposing directions (Figure 2F).

The addition of cholesterol to cells increased the mitochondrial membrane lipid order of HAECs under static conditions, and it markedly suppressed the laminar flow-induced decrease in the lipid order (Figure 2D, F). On the other hand, cholesterol extraction by treating the cells with MβCD significantly decreased the mitochondrial membrane lipid order of cells under static conditions, and it clearly prevented the disturbed flow-induced increase in the lipid order (Figure 2E, F). These findings indicate that adding or removing cholesterol to cells affects the lipid order of mitochondrial membranes and can prevent the effects of laminar and disturbed flow on the mitochondrial membrane lipid order.

### Laminar flow rapidly increased mitochondrial ATP production, whereas disturbed flow decreased ATP production

HAECs were exposed to laminar flow or disturbed flow, and changes in the mitochondrial ATP levels were examined using real-time imaging with mitAT1.03, a genetically encoded FRET-based ATP biosensor that targets the mitochondrial matrix.^23^ Pseudocolor images of the YFP/CFP ratio showed increased ATP levels over the entire mitochondria when the cell was exposed to laminar flow (Figure 3A, F). In contrast, cells subjected to disturbed flow exhibited decreased ATP levels (Figure 3B, F). The temporal changes in ATP levels were quantified by defining ROIs throughout the cell. The ATP level quickly increased in response to laminar flow but decreased in response to disturbed flow, and both the increased and decreased ATP levels returned to their original levels after the flows ceased. Similar reactions occurred in response to a second flow loading, indicating that the mitochondrial responses to laminar and disturbed flows were reversible.

**Figure 3.**
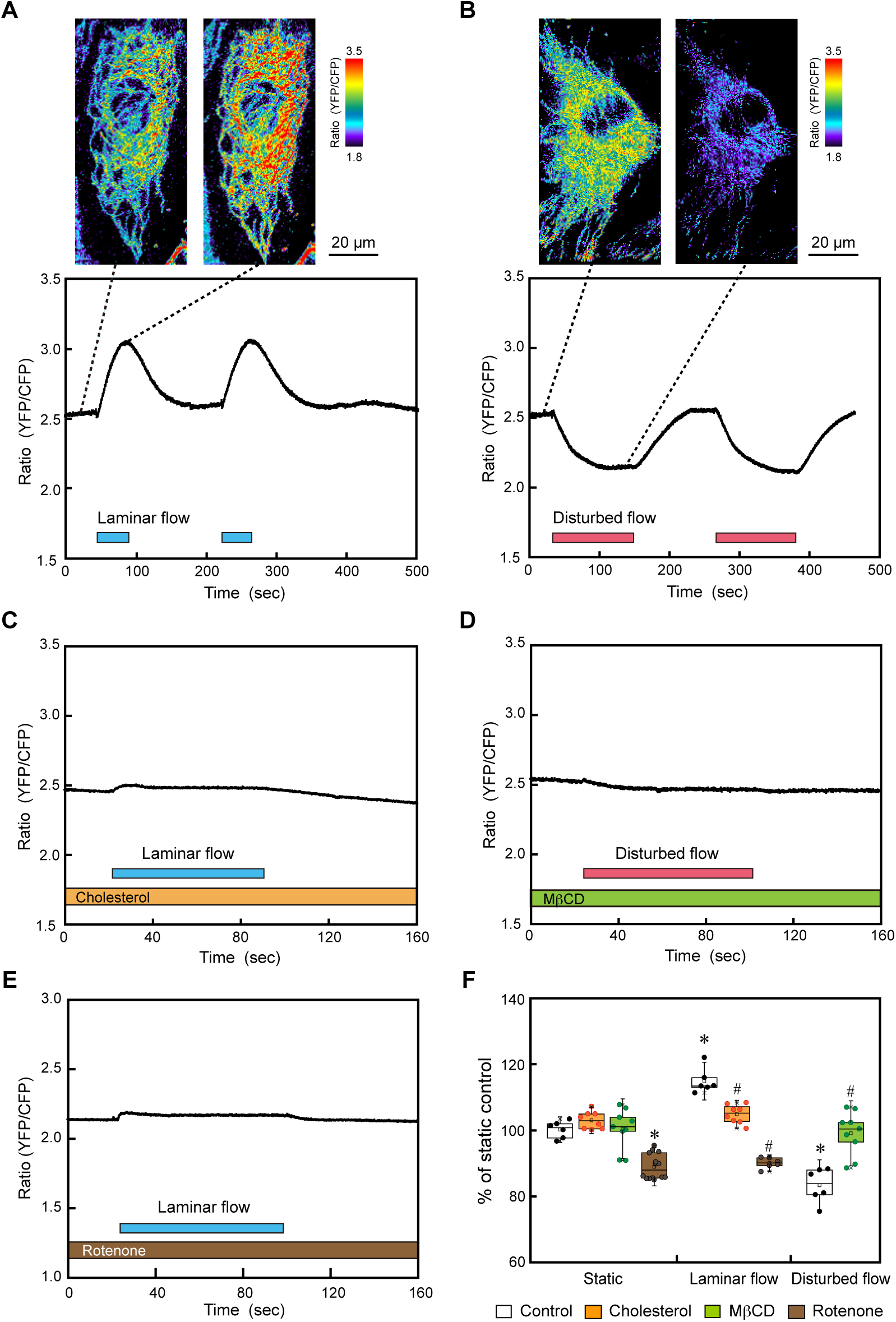
Laminar versus disturbed flow effects on mitochondrial ATP production. HAECs were exposed to laminar flow or disturbed flow and changes in mitochondrial ATP production were examined using real-time imaging with the mitochondria-targeting ATP biosensor mitAT1.03. (**A**) Mitochondrial ATP responses to laminar flow. Pseudocolor images of the mitAT1.03 fluorescence ratio (YFP/CFP) exhibited an increase in ATP levels when the cell was exposed to laminar flow. The temporal changes in the mitAT 1.03 ratio showed that two similar responses of increasing ATP levels occurred for two repeated laminar flow loadings. (**B**) Mitochondrial ATP responses to disturbed flow. Pseudocolor images exhibited a decrease in ATP levels when the cell was exposed to disturbed flow. Temporal changes showed that two similar responses of decreasing ATP levels occurred for two repeated disturbed flow loadings. (**C**) Addition of cholesterol (100 μM) to ECs markedly inhibited the laminar flow-induced increase in mitochondrial ATP production. (**D**) Cholesterol extraction by treating ECs with MβCD (10 mM) abolished the disturbed flow-induced decrease in mitochondrial ATP production. (**E**) Treatment of ECs with rotenone (5 μM), an inhibitor of the mitochondrial electron transport system, lowered the mitochondrial ATP levels in cells under static conditions and abolished the ATP increasing effect of laminar flow. (**F**) Quantitative analysis of the changes in mitochondrial ATP production induced by laminar flow and disturbed flow, with or without cholesterol, MβCD, or rotenone. Values are the means ± SD of the data obtained from samples shown as points. \**P* < 0.01 vs. control. ^#^*P* < 0.01 vs. each flow control.

The addition of cholesterol to the cells abolished the laminar flow-induced increase in mitochondrial ATP levels (Figure 3C, F), and the treatment of cells with MβCD prevented the disturbed flow-induced decrease in mitochondrial ATP levels (Figure 3D, F). These findings indicate that changes in membrane physical properties, such as lipid order and cholesterol content, are closely linked to the effects of laminar and turbulent flows on mitochondrial ATP production.

The treatment of cells with rotenone, an inhibitor of mitochondrial electron transport chain complex I, abolished the laminar-flow-induced increase in the ATP level, indicating that the changes in ATP levels were due to changes in mitochondrial oxidative phosphorylation (Figure 3E, F).

### Laminar flow decreased mitochondrial H_2_O_2_ release, whereas disturbed flow increased H_2_O_2_ release

HAECs were exposed to laminar flow or disturbed flow, and changes in mitochondrial H_2_O_2_ release were examined using real-time imaging with HyPer7, a genetically encoded high-affinity H_2_O_2_ sensor that targets the mitochondrial matrix.^24^ Pseudocolor images of the HyPer7 fluorescence intensity ratio (F500/F438) exhibited a clear decrease in H_2_O_2_ levels when the cell was exposed to laminar flow (Figure 4A). The temporal changes in the H_2_O_2_ level were quantified by setting ROIs throughout the cell, and three similar reductions in H_2_O_2_ levels were observed for three repeated laminar flow loadings. In contrast, pseudocolor images exhibited a clear increase in H_2_O_2_ levels when the cells were exposed to disturbed flow (Figure 4B). The temporal changes showed three similar increases in H_2_O_2_ levels in response to three repeated disturbed flow loadings. Both the laminar flow-induced decrease and the disturbed flow-induced increase in H_2_O_2_ levels returned to the control level after each flow ceased, indicating that the mitochondrial H_2_O_2_ releasing responses were reversible. A quantitative analysis of several cells confirmed that mitochondrial H_2_O_2_ release exhibited contrasting responses, decreasing with laminar flow and increasing with disturbed flow (Figure 4F).

**Figure 4.**
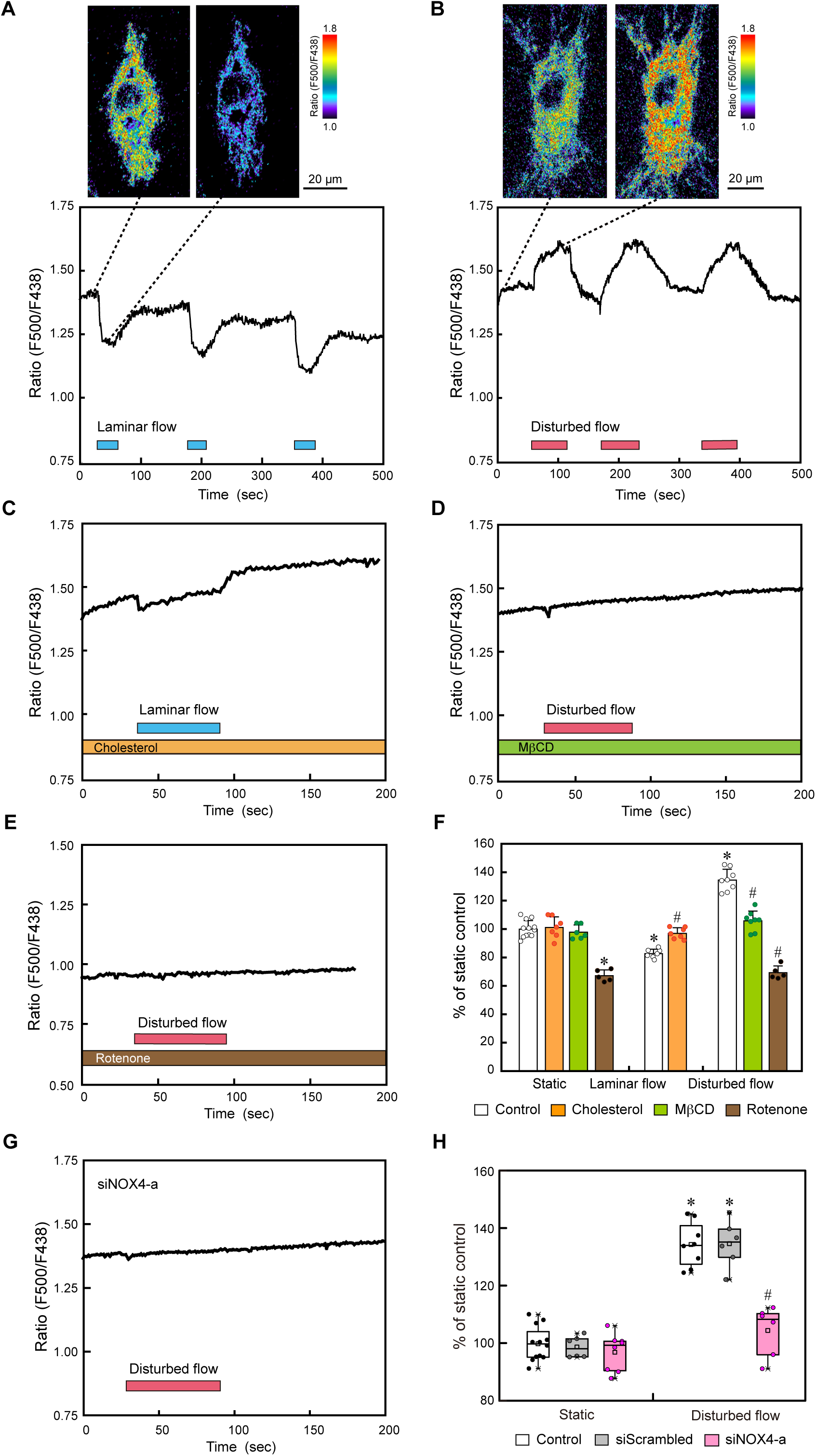
Laminar versus disturbed flow effects on mitochondrial H_2_O_2_ release. HAECs were exposed to laminar flow or disturbed flow, and changes in mitochondrial H_2_O_2_ release were examined using real-time imaging with a mitochondrial matrix-targeted H_2_O_2_ sensor probe, HyPer7. (A) Mitochondrial H_2_O_2_ responses to laminar flow. Pseudocolor images of the HyPer7 fluorescence intensity ratio (F500/F438) exhibited a decrease in H_2_O_2_ levels when the cell was exposed to laminar flow. The temporal changes in the HyPer7 ratio showed that three similar responses of decreasing H_2_O_2_ levels occurred for three repeated laminar flow loadings. (B) Mitochondrial H_2_O_2_ responses to disturbed flow. Pseudocolor images exhibited an increase in H_2_O_2_ levels when the cell was exposed to disturbed flow. The temporal changes showed that three similar responses of increasing H_2_O_2_ levels occurred for three repeated disturbed flow loadings. (C) Addition of cholesterol (100μM) to ECs markedly inhibited the laminar flow-induced decrease in mitochondrial H_2_O_2_ release. (D) Cholesterol extraction by treating ECs with MβCD (10mM) abolished the disturbed flow-induced increase in mitochondrial H_2_O_2_ release. (E) Treatment of ECs with rotenone (5 μM) abolished the disturbed-flow-induced H_2_O_2_ increase. (F) Quantitative analysis of the changes in mitochondrial H_2_O_2_ release induced by laminar flow and disturbed flow, with or without cholesterol, MβCD, or rotenone. Values are the means ± SD of the data obtained from the samples shown as points. \**P* < 0.01 vs. control. ^#^*P* < 0.01 vs. each flow control. (G) Knockdown of Nox4 expression with siNOX4-a blocked the disturbed flow-induced H_2_O_2_ increase. (H) Quantitative analysis of the changes in mitochondrial H_2_O_2_ release induced by disturbed flow, with or without siScrambled or siNOX4-a. Values are the means ± SD of the data obtained from the samples shown as pointes. \**P* < 0.01 vs. control. ^#^*P* < 0.01 vs. each flow control.

The addition of cholesterol to ECs markedly suppressed the laminar flow-induced decrease in mitochondrial H_2_O_2_ release (Figure 4C, F). On the other hand, cholesterol extraction by treating ECs with MβCD abolished the disturbed flow-induced increase in mitochondrial H_2_O_2_ release (Figure 4D, F). These findings indicate that changes in the membrane physical properties, such as lipid order and cholesterol content, are closely linked with the effects of laminar and disturbed flows on mitochondrial H_2_O_2_ release.

Furthermore, when ECs were treated with rotenone, disturbed flow did not increase the H_2_O_2_ levels in ECs, indicating that the mitochondrial electron transfer system was involved in the disturbed flow-induced increase in H_2_O_2_ levels (Figure 4E, F).

Next, we examined whether NADPH oxidase (Nox) was involved in the increase in H_2_O_2_ levels. Real-time PCR showed that the Nox isoform 4 (Nox4) was predominantly expressed at the mRNA level and that other isoforms, including Nox1, 2, 3, and 5, were barely expressed in the HAECs used in this study (Figure S1A). Immunostaining with a Nox4 antibody showed that the Nox4 protein was predominantly localized to mitochondria in HAECs (Figure S1B), which was consistent with the findings of a previous report.^25^ A small-interfering RNA (siRNA), siNOX4-a, effectively knocked down the gene expression of Nox4 in HAECs, thereby abolishing the disturbed flow-induced increase in H_2_O_2_ levels (Figure S1C, Figure 4G, H). Taken together, these findings indicate that the increased H_2_O_2_ originated from reactive oxygen species (ROS) produced through the cooperation of the mitochondrial electron transport system and Nox4.

## Discussion

A variety of *in vitro* models have been used to study the effect of disturbed flow on ECs.^3^ These include disturbed flow-loading systems using parallel-plate flow chambers or cone-and-plate viscometers. Among these models, the flow chamber system has the advantage of providing a simple, predictable, and controllable disturbed flow to the cells. In this study, we used a flow system in which a step is placed upstream of the flow path of the parallel-plate flow chamber to generate disturbed flow. This system allows direct microscopic observation of the cells under flow conditions and real-time imaging with various photoaffinity probes. More importantly, the system was able to reproduce the desired *in vivo* disturbed flow by adjusting the height of the step based on a CFD analysis of medium flow in the chamber. As a hemodynamic metric that quantitatively indicates the multidirectional character of disturbed flow, we used the transverse wall shear stress (transWSS), which is calculated as the time-average of wall shear stress components perpendicular to the mean flow direction. TransWSS was shown to be an important factor in determining the site of origin of diet-induced plaques at intercostal artery bifurcation orifices in rabbits,^26^ and the normalized transWSS (NtransWSS), which corresponds to transWSS divided by the time-averaged WSS, was correlated most strongly with the location and size of human cerebral aneurysms among various hemodynamic metrics, including the time-averaged WSS gradient, oscillatory shear index, and gradient oscillatory number.^27^

The present study showed that the plasma membrane immediately responds to disturbed flow by increasing their lipid order. This was the opposite of the response to laminar flow, in which the lipid order was reduced, indicating that the plasma membrane itself can differentiate flow patterns between laminar and disturbed flow. Changes in the physical properties of the plasma membrane, such as lipid order, fluidity, and viscosity, are known to have a direct effect on the conformation and dynamics of lipid and protein molecules present in the plasma membrane, modifying their function.^28, 29^ Numerous studies have shown that flow stimuli to ECs activate a variety of membrane molecules, including ion channels such as Piezo-1 and TrpV4, receptors such as G protein coupling receptors and vascular endothelial growth factor receptors (VEGFRs), and adhesion molecules such as vascular endothelium cadherin and platelet-EC adhesion molecule-1.^9, 30–34^ Accordingly, it is highly possible that changes in the lipid order of plasma membranes are involved in the flow-induced activation of many, if not all, of the above-mentioned membrane molecules. Taken together, ECs seem to have a flow-sensing and signaling mechanism in which their plasma membranes act as a sensor to separate laminar and disturbed flow, and membrane-associated molecules, microdomains and subcellular organelles act as transducers to transmit information downstream.

The current study demonstrated that flow stimuli to ECs changed the lipid order of not only plasma membranes, but also mitochondrial membranes. Similar to the effects on plasma membranes, laminar flow decreased the lipid order of the mitochondrial membranes, while disturbed flow increased the lipid order. How such responses occur in mitochondrial membranes, which are not directly exposed to flow, remains unknown. Several possible mechanisms can be postulated. Flow-induced changes in the lipid composition of the plasma membrane, such as changes in cholesterol content, may lead to similar changes in the mitochondrial membrane. In fact, cell treatments involving the addition or removal of cholesterol blocked the laminar and disturbed flow-induced changes in lipid order in both plasma and mitochondrial membranes. Another mechanism is that since mitochondria are bound to the cytoskeleton (for example, to actin filaments), mechanical stress transmitted through the cytoskeleton may act on mitochondrial membranes to alter their lipid order.^35, 36^ Alternatively, when fluid shear stress was applied to liposomes that were made of artificial lipid bilayers, flow was reportedly generated within the liposomes.^37, 38^ Although the mechanism responsible for this phenomenon is unclear, shear stress may induce the flow of liquid components within ECs, directly affecting mitochondrial membranes. Understanding this process will require further research.

In ECs, the production of ATP as energy is mainly dependent on the glycolytic system, and mitochondria-produced ATP is thought to act as a signaling molecul.^15^ Our previous study showed that mitochondria play a critical role in EC signaling of laminar flow by increasing ATP production, leading to purinergic Ca^2+^ signaling^20^; the present study revealed that mitochondria are also involved in the signaling of disturbed flow. Real-time imaging with Hyper7 demonstrated that disturbed flow caused an immediate increase in mitochondrial H_2_O_2_ release. This response was abolished by the treatment of cells with rotenone, an electron transfer system inhibitor. Furthermore, Nox4, which catalyzes the production of a superoxide free radical, was shown to be involved in the disturbed flow-induced H_2_O_2_ release. Thus, the increased H_2_O_2_ was thought to originate from ROS produced by the cooperation of the mitochondrial electron transport system and Nox4. Thus, EC mitochondria appeared to transmit information regarding disturbed flow downstream via ROS signaling.

In the current study, we found that changes in the physical properties of plasma and mitochondrial membranes were closely linked to changes in mitochondrial functions, such as ATP and ROS production. The blockage of flow-induced changes in the lipid order of both membranes by the addition or removal of cholesterol abolished the laminar flow- or disturbed flow-induced ATP production or H_2_O_2_ release, respectively. Concerning how changes in the lipid order of mitochondrial membranes affect oxidative phosphorylation to produce ATP or ROS, several possible mechanisms were proposed: 1) changing the permeability of the membrane to oxygen and other substrates necessary for respiration, 2) affecting the function of enzymes working in electron transport systems, and 3) influencing the assembly and function of respiratory chain complexes by altering the diffusion of molecules within the membrane.^21^

To date, many studies have shown that oscillatory flow, a pattern of disturbed flow, increases ROS production in ECs.^39–43^ When cultured ECs, such as human umbilical ECs, murine aortic ECs, and bovine aortic ECs, were exposed to oscillatory flow with a velocity that oscillated back and forth with a periodicity of ±3∼5 dyne/cm^2^, 1 Hz, the production of O_2_^-^ and H_2_O_2_ significantly increased from 30 minutes to 24 hours later. NADPH oxidases, including Nox1, Nox2 and Nox4, were also shown to be involved in oscillatory-flow-induced ROS production, although the isoform of Nox involved differed depending on the cell type. The present study showed that the expression of Nox4 was prominent in HAECs, but Nox1, Nox2, and Nox5 were barely expressed; in addition, Nox4-mediated ROS production contributed considerably to the disturbed flow-induced increase in mitochondrial H_2_O_2_ release. Recently, Nox4 was shown to be localized to the mitochondrial inner membrane in human renal epithelial cells and to have an ATP-binding motif; its superoxide generation activity was also found to be negatively regulated by mitochondrial ATP levels.^44^ Thus, this study showed that disturbed flow had the effect of lowering the mitochondrial ATP levels, which may have activated Nox4 to increase ROS production within the mitochondrial compartment.

Mitochondria-produced ROS were once thought to be harmful byproducts that impaired both mitochondria and cellular functions. Recently, however, mitochondrial ROS have been shown to be deeply involved in the maintenance of cellular homeostasis and adaptations to environmental stresses.^45–47^ Among ROS, H_2_O_2_ is highly stable, has a long half-life *in vivo*, and can pass freely through cell membranes, as it has no electrical charge. When released from ECs, H_2_O_2_ acts as an endothelial hyperpolarizing factor in smooth muscles and dilates blood vessels.^48^ H_2_O_2_ also plays a role in angiogenesis and vascular remodeling by augmenting EC migration and proliferation via the phosphorylation of VEGFRs.^49, 50^ In addition, H_2_O_2_ mediates the expression of heme oxygenase-1,^51^ which protects cells from inflammation, and the stabilization of hypoxia-inducible factor-1, which is required for the adaptive response to hypoxia. ROS also cause reversible post-translational modifications of proteins via oxidation and regulate several signaling pathways.^52^ Therefore, H_2_O_2_ released by mitochondria is thought to act as a second messenger to inform cells of disturbed flow and to trigger cellular responses that allow adaptation via redox signaling. However, the specific molecular targets of mitochondrial H_2_O_2_ produced in response to disturbed flow are currently unknown, and identifying these targets will be an important topic for future research. On the other hand, as a long-term effect of disturbed flow, if ECs are exposed to cardiovascular risk factors such as hypertension, dyslipidemia, and hyperglycemia, and if mitochondria cooperate with other intracellular sources of ROS to produce excessive ROS, EC dysfunction and the activation of proinflammatory pathways would occur; if sustained, this would lead to the development of atherosclerotic plaques and aneurysms.^46^

In summary, the current study demonstrated that ECs perform distinct sensing and signaling for laminar and disturbed flows, in which both plasma membranes and mitochondrial membranes play critical roles by altering their lipid order in opposite directions, realizing intracellular signaling specific to each flow pattern. The changes in membrane lipid order were closely linked to changes in mitochondrial functions, i.e., increased ATP production for laminar flow and increased H_2_O_2_ release for disturbed flow. Thus, laminar flow information seems to be transmitted downstream as ATP-mediated purinergic signaling, while disturbed flow information is transmitted as ROS-mediated redox signaling. However, many unsolved problems remain, such as the biophysical or thermodynamic mechanisms by which shear stress alters the plasma membrane lipid order, the mechanism by which shear stress acting on the plasma membrane affects the mitochondrial membrane properties, and the mechanism by which changes in the mitochondrial membrane lipid order modulate the electron transfer system and Nox to produce ATP and ROS. Resolving these problems in the future will contribute not only to a better understanding of EC sensing and signaling mechanisms for different blood flow patterns, but also to the elucidation of their roles in the maintenance of vascular physiology and in pathological conditions, such as hypertension, thrombosis, aneurysms, and atherosclerosis. It is also expected to contribute to the development of new preventive and therapeutic strategies for these vascular diseases.

## Materials and Methods

### Cell cultures and treatments

HAECs were purchased from Lonza Group Ltd. (Material number: cc-2535) and were grown in M199 supplemented with 15% fetal bovine serum, 2 mM L-glutamine (Gibco), 50 µg/mL heparin, and 30 µg/mL EC growth factor (Becton Dickinson) in a 1% gelatin-coated tissue culture flask. The cells used in the experiments in this study were in the 7th and 10th passages. All the experiments were approved by the Ethics Committee of the University of Tokyo, Graduate School of Medicine.

Cholesterol reduction was induced by incubating HAECs with 10 mM methyl-β-cyclodextrin (MβCD, Sigma-Aldrich) dissolved in Hanks’ Balanced Salt solution (HBSS, Sigma-Aldrich) for 30 min at 37°C. Cholesterol enrichment was conditioned by incubating HAECs with a complex of cholesterol (Sigma-Aldrich) and MβCD (1:7 molar ratio) dissolved in the complete culture medium for 6 hours at 37°C. The final concentration of cholesterol was 100 μM. For the inhibition of mitochondrial respiratory chain complex I, HAECs were treated with 5 μM rotenone (Sigma-Aldrich) dissolved in HBSS for 10 min at 37°C.

### Flow-loading experiments

We fabricated a parallel-plate dynamic flow system that produces disturbed flow mimicking the flow characteristics in human arteries. The flow chamber consisted of a 1%-gelatin-coated cover glass with the cultured cells on one side and a second, parallel, glass plate held 700 μm apart from the first plate by a Teflon^TM^ gasket. The medium was perfused using a roller/tube pump, and the entire closed circuit was maintained at exactly 37°C. The flow chamber had a motorized height-changeable step near the inlet, which created a disturbed flow characterized by a recirculation eddy immediately downstream of the step, followed by a region of flow reattachment. As a metric for disturbed flow, we used the normalized transverse wall shear stress (NtransWSS), which was calculated using a computer fluid dynamic (CFD) analysis based on the geometrical parameters of the flow chamber and the viscosity, density, and velocity of the culture medium.^27^ NtransWSS represents the degree of disturbance in the strength and direction of the shear stress, with a maximum value of 1.0. Since disturbed flow with an NtransWSS ranging from 0.2 to 1.0 has been shown to occur in human carotid bifurcations where atherosclerotic plaques are prone to develop,^53^ we subjected the cells to a disturbed flow with an NtransWSS of 0.2 or more (average, 0.4) and an average WSS of 20 dynes/cm^2^ by adjusting the step height and flow rate.

On the other hand, a parallel-plate flow chamber without steps was used for experiments in which laminar flow was applied to ECs. The intensity of the shear stress (τ, dynes/cm^2^) acting on the EC layer was calculated from the formula τ = 6μQ/*a*^2^*b*, where μ is the viscosity of the perfusate (poise), Q is the flow volume (mL/s), and *a* and *b* are the cross-sectional dimensions of the flow path (cm). In the present experiments, the cells were exposed to laminar flow with an NtransWSS of zero and a WSS of 15 dyne/cm^2^, which was within the range of physiological levels (10-20 dynes/cm^2^) in large to medium-sized human arteries.^54^

### Computational Fluid Dynamics (CFD) analysis

To analyze the fluid flow and quantify NtransWSS in the flow-loading device, we first generated computational meshes with a resolution of 0.02-0.05 mm and a total element number of 9,268,480 for the geometry model of the device using the Mixed-Element Grid Generator in 3 Dimensions (MEGG3D).^55^ The fluid flow in the device was simulated by solving the Navier-Stokes equations using the CFD software OpenFOAM 8 (OpenCFD Ltd.); NtransWSS was then quantified on the bottom of the device. In this simulation, the HBSS fluid was treated as an incompressible Newtonian viscous fluid with a density of 1,010 kg/m^3^ and a viscosity of 0.77 mPa·s, and its flow rate was set to 200 mL/min.

### Laurdan imaging

Real-time Laurdan imaging was performed to analyze the lipid order of the plasma membranes, as described previously.^11^ Lipid order is a physical property of lipid bilayer membranes and depends on the type, distribution, and density of lipids, the orientation and kinetic state of the acyl chains of phospholipids, and the cholesterol content. As the lipid order increases, the cell membrane becomes more rigid and less fluid. Cells labeled with Laurdan dye (Molecular Probes) were placed in mechanical force-loading devices and then mounted on the stage of a confocal laser-scanning microscope (TCS SP2 AOBS, Leica Microsystems) equipped with a two-photon laser (MaiTai BB, Spectra-Physics) and HCX-PL APO 100x/1.40-0.70 oil objective. Laurdan fluorescence was excited with a 770-nm wavelength light, and the emitted light was collected in the 400-460 nm range for one channel and the 470-530 nm range for the other channel. General polarization (GP) was calculated using the following formula, as previously described:^56^

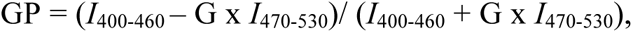

where *I* is light intensity and G is a correction factor calculated from two variables: a known GP value for Laurdan in DMSO at 22°C, and the GP value of the Laurdan stock solution (500 μM) determined in each experiment. GP images (512 x 512 pixels) were analyzed and pseudocolored with ImageJ1.44n software (NIH Image). The GP values, which reflect the extent of water penetration into a lipid bilayer, were used as an indicator of the membrane lipid order.

### NRMito imaging

The lipid order of mitochondrial membranes was examined using real-time imaging with the solvatochromic fluorescent probe NRMito (a kind gift from Dr. Andrew S. Klymchenko), a Nile Red derivative bearing chemical groups for the specific targeting of mitochondria.^22^ HAECs were incubated with 20 nM NRMito for 45 min at 37°C. The localization of NRMito in the mitochondria of HAECs was detected using 50 nM of MitoTracker Green FM (Invitrogen). The NRMito imaging was performed using a confocal laser-scanning microscope (TCS SP2 AOBS, Leica Microsystems) with an HCX-PL APO 63x/1.40-0.60 oil objective and argon laser at 37°C. NRMito fluorescence was excited with light at a 514-nm wavelength, and the emitted light was collected in the 550-600 nm range for one channel and the 600-650 nm range for the other channel. The ratiometric images were analyzed using MetaMorph software, version 7.7 (Molecular Devices).

### Imaging of Mitochondrial ATP

The mitochondrial ATP was imaged using a FRET-based ATP biosensor targeted at the mitochondrial matrix (mitAT1.03; a kind gift from Dr. Hiromi Imamura), as previously described.^23^ The cDNA of mitAT1.03 was ligated and cloned into the *Sal*I/*Not*I sites of a pAd-CMV-V5-DEST Gateway Vector (Thermo Fisher Scientific), and the adenovirus vector containing mitAT1.03 cDNA (Ad-mitAT1.03) was constructed as described previously. Cultured HAECs were infected with 50 pfu/cell of Ad-mitAT1.03. Three to five days after Ad-mitAT1.03 infection, the HAECs (maintained at 37°C) were imaged using an ECLIPSE Ti-E inverted microscope (Nikon) with 100× Apo TIRF oil objective (NA = 1.49) using a water-cooling electron multiplier CCD camera (ImagEM C9100-13; Hamamatsu), controlled by HCImage software, version 4.3.5.6 (Hamamatsu). Dual-emission ratio imaging of mitAT1.03 was performed using an FF01-427/10 excitation filter (Semrock), an FF458-Di01 dichroic mirror, and a Dual View Multichannel Imaging System (DV2, Photometrics) equipped with two emission filters (FF01-483/32 for CFP and FF01-542/27 for YFP). The images were analyzed using MetaMorph software, version 7.7 (Molecular Devices).

### Imaging of mitochondrial H_2_O_2_ release

Mitochondrial ROS were imaged using a mitochondria matrix targeted H_2_O_2_ probe, HyPer7.^24^ The cDNA of pCS2+MLS-HyPer7 was purchased from Addgene (Plasmid ID: 136470) and was transfected in HAECs using the Neon^TM^ Transfection System (Invitrogen) according to the manufacturer’s recommendations. Three days after transfection, real-time imaging was performed using an ECLIPSE Ti-E inverted microscope (Nikon) with 40× Plan Apo silicon oil λS objective (1.25 numerical aperture [NA]) using a water-cooling electron multiplier CCD camera (ImagEM C9100-13; Hamamatsu), controlled by NIS Element software, version 4.30 (Nikon). The fluorescence of HyPer7 probes was excited sequentially via FF02-438/24 and FF01-500/24 band-pass excitation filters (Semrock). Emissions were collected every 0.1 s using a FF01-542/27 band-pass emission filter cube equipped with a FF520-Di02-25x36 dichroic mirror (Semrock). Images of the HyPer7 fluorescence intensity ratio (F500/F438) were analyzed using MetaMorph software, version 7.7 (Molecular Devices).

### Statistical Analysis

The results were presented as the mean ± S.D. Statistical significance was evaluated using an ANOVA with the Bonferroni adjustment applied to the results of post-hoc *t* tests. The statistical analysis was performed using SPSS software (SPSS Inc.). A *P* value of <0.01 was regarded as being statistically significant.

## Acknowledgments

We wish to acknowledge the following people for providing materials or technical assistance: Dr. Dmytro I. Danylchuk and Dr. Andrew S. Klymchenko (University of Strasbourg) for providing the NRMito, Dr. Hiromi Imamura for providing the ATeam plasmid vector, and Ms. Yuko Sawada for her technical assistance, including the plasmid construction and cell cultures.

## Author contributions

Both K.Y. and J.A. designed the study, conducted the experiments, analyzed the results, and wrote the manuscript. Y.S. performed the CFD analysis. R.M. and K.K. analyzed the data.

---

**Figure S1.**
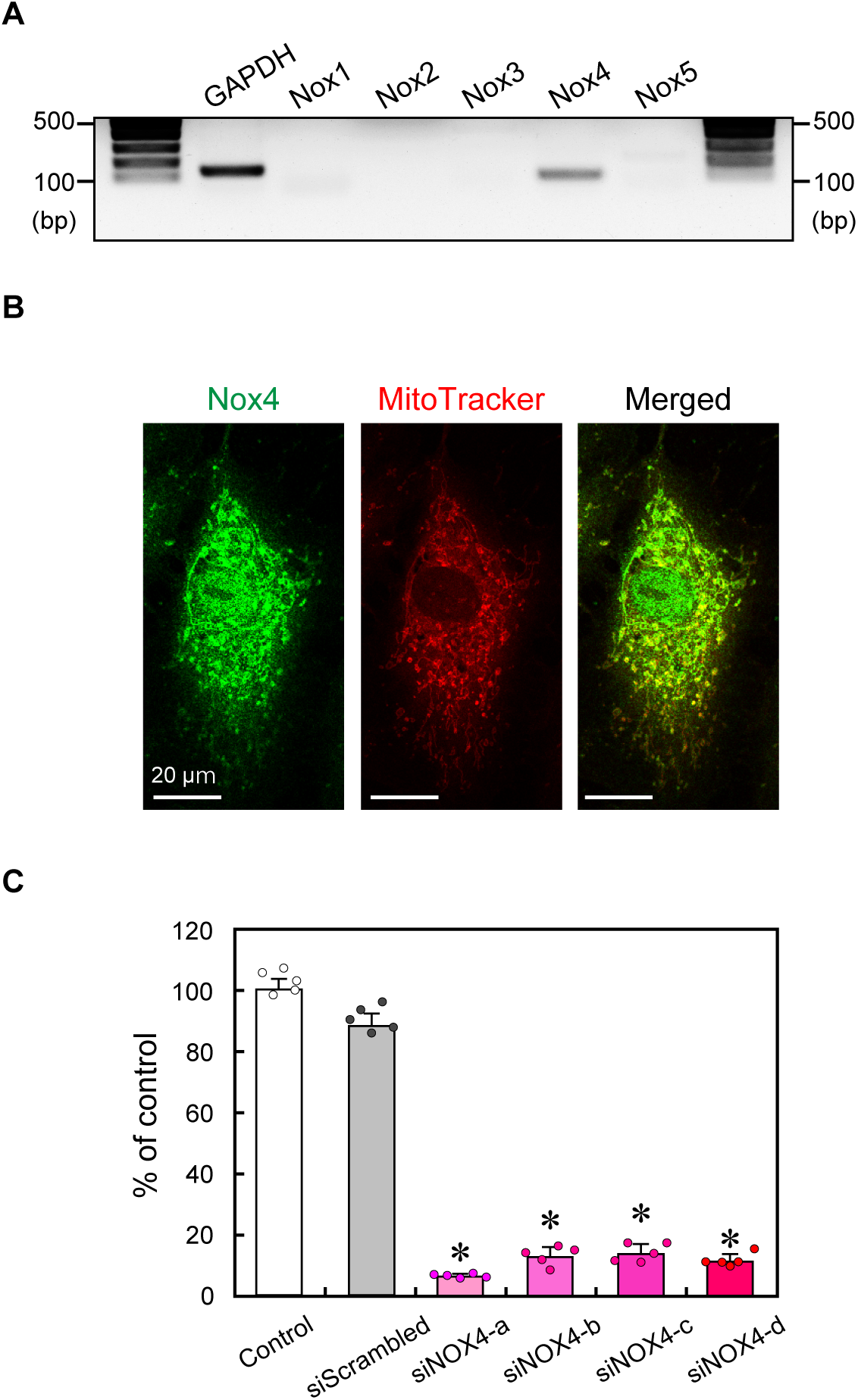
(A) The mRNA expression of Nox isoforms was determined using RT-PCR. Nox4 was predominantly expressed, but Nox1, 2, 3, and 5 were barely expressed in HAECs. (B) Confocal microscopic images of HAECs were stained with anti-Nox4 antibody (in green) and labeled with the mitochondrion selective dye MitoTracker Deep Red (in red). The majority of Nox4 was localized to the mitochondria. (C) Quantitative mRNA expression of Nox4 determined by RT-qPCR. All four siRNAs (siNOX4-a, b, c, d) effectively suppressed the mRNA expression of Nox4. Values are the means ± SD of the data obtained from samples shown as points.

